# Abscisic Acid rescues behavior in adult female mice in Attention Deficit Disorder with Hyperactivity model of dopamine depletion by regulating microglia and vesicular GABA transporter

**DOI:** 10.1101/2024.05.15.592910

**Authors:** Maria Meseguer-Beltrán, Sandra Sánchez-Sarasúa, Nóra Kerekes, Marc Landry, Matías Real-López, Ana María Sánchez-Pérez

**Author notes:** To whom correspondence may be addressed: Ana María Sánchez-Pérez.

## Abstract

**Background:** Attention deficit/hyperactivity disorder (ADHD) is a neurodevelopmental syndrome influenced by both genetic and environmental factors. While genetic studies have highlighted catecholamine dysfunction, emerging epidemiological evidence suggest neuroinflammation as a significant trigger. However, understanding the relative contributions of these alterations to ADHD symptomatology remains elusive.

**Method:** This study employed 93 female Swiss mice of the ADHD dopamine deficit model. Dopaminergic lesions were induced via 6-hydroxidopamine (6-OHDA) injection on postnatal day 5. The impact of these lesions during development was examined by comparing young and adult mice (at postnatal day 21 and 90, respectively). We sought to mitigate adult symptoms through abscisic acid (ABA) administration during two-months. Postmortem analyses encompassed the evaluation of neuroinflammation (microglia morphology, NLRP3 inflammasome activation, cytokine expression) and excitatory/inhibitory (E/I) ratio in specific brain regions.

**Results:** Neonatal dopaminergic lesions elicited hyperactivity, impulsivity, hypersensitivity increased social interaction in both one-month and three-month females and induced impaired memory in three-month mice. ABA exposure significantly ameliorated hyperactivity, impulsivity, anxiety, hypersensitivity, and social interaction alterations, but not cognitive impairment. In the anterior cingulate cortex (ACC) of one-month mice dopamine-deficit elevated IL-1β and TNFα expression and reduced Arg1 mRNA levels, along with E/I imbalance. ABA intervention restored microglia morphology, IL-1β, Arg1 expression and enhanced vGAT levels.

**Conclusions:** This study strongly suggest that dopamine deficit induced alteration of microglia and E/I ratio underling distinct ADHD symptoms. Reinstating healthy microglia by anti-inflammatory agents in specific areas emerges as a promising strategy for managing ADHD.

## Introduction

Attention deficit/hyperactivity disorder (ADHD) is a neurodevelopmental disorder that affects 5 to 10% of children and can persist into adulthood. It is estimated to affect up to 2.5% of adults worldwide (1). ADHD profoundly impacts patients’ functioning and quality of life (2–4), thereby increasing the risk of educational and occupational failure. Suffering from ADHD is associated with higher social disability, criminality, and addictions if not properly managed (5–7). The diagnosis of ADHD includes the occurrence of hyperactivity, impulsivity, and sustained inattention. However, these three symptoms may not occur simultaneously, and patients may present as inattention-type or impulsive-hyperactive type. Additionally, ADHD core symptoms may co-occur with other psychiatric traits, such as anxiety (8), depression (9), and somatic complaints, including hypersensitivity to mechanical or thermal stimulus (10), indicating a complex etiology at the onset and progression of the disease.

ADHD onset has been associated to catecholamine dysfunction, and consequently, medications that increase dopamine (DA) and norepinephrine signaling by blocking the transporters are often prescribed (11). However, chronic prescription of these drugs to children may result in tolerance, dependence/addiction (12). In addition, adverse effects have been described including cardiovascular risks and depression (13). At the neurobiological level, alterations in the anterior cingulate cortex (ACC) are linked to the occurrence of ADHD in humans and rodent models (14,15). The ACC is a hub cortical area that receives sensitive information via the thalamus and DA terminals from the ventral tegmental area (VTA) (16), the main source of DA in the brain. In turn, the ACC projects to other cortical areas, thereby modulating a wide variety of behavioral traits that include core symptoms of ADHD like inattention, impulsivity and hyperactivity (17,18). It also modulates pain perception, via descending projections to the posterior insula (pIC), (for review see (19)); and anxious behavior via connections with the amygdala (20,21). Thus, ACC hyperexcitability is associated with increased pain (22) and anxious behavior (23). The ACC is a hub controlling pain and pain related emotions (24,25).

There is increasing evidence pointing to the influence of inflammation during early development as a risk factor aggravating and/or triggering the symptoms (26–29). In this line, microbial dysbiosis (30), and maternal autoimmune reactions (31) have been strongly associated with ADHD incidence and co-existing conditions (32).

Microglia cells, contribute significantly to the modeling and maintenance of neural networks (33,34), via synapse elimination and release of neurotrophic factors. In neuroinflammation with overactivated microglia synaptic elimination is disturbed leading to a disruption of the excitatory/inhibitory (E/I) balance and hyperexcitability (35). E/I imbalance underlies cognitive impairment and altered social and emotional behaviors (36,37), contributing to the etiology of neurological, mental and developmental diseases (38). In addition to induce an E/I imbalance, activated microglia release proinflammatory cytokines (i.e. IL-1β/IL-18) via NLRP3 inflammasome signaling (39–41), or tumor necrosis factor α (TNFα) (42); reactive oxygen species, glutamate, amongst other biologically active substances. To terminate the inflammatory process, active microglia switch from the M1 to the M2 status, with a distinct expression profile. The enzyme Arginase 1 (Arg1), which competes with iNOS to reduce nitric oxide production (43,44) is characteristic of M2 status.

We have previously observed microglia alterations in the ACC and pIC in females and males of a validated mice model of ADHD, the neonatal 6-hydroxydopamine (6-OHDA) lesion (45), that were rescued by one-month of abscisic acid (ABA) treatment, alleviating pain sensitivity in female mice and hyperactivity in males (46). ABA is an evolutionary well-conserved hormone synthesized endogenously in a wide spectrum of cells (47). ABA via PPARγ signaling, has been shown to inhibit NLRP3 inflammasome activation and oxidative stress (48). Therefore, we hypothesized that long-term effects of ABA treatment in dopamine deficit model of ADHD would rescue symptomatology via microglia, NLRP3 inflammasome and E/I modulations in specific brain areas. Furthermore, as age is an important factor modulating ADHD symptoms, we compared behavior at postnatal day 21 (P21), and P90. Three weeks and three months in mice is considered equivalent to human infant and young adults, respectively (49).

## Methods and Materials

The time course of experiments is described in Fig. 1.

**Figure 1.**
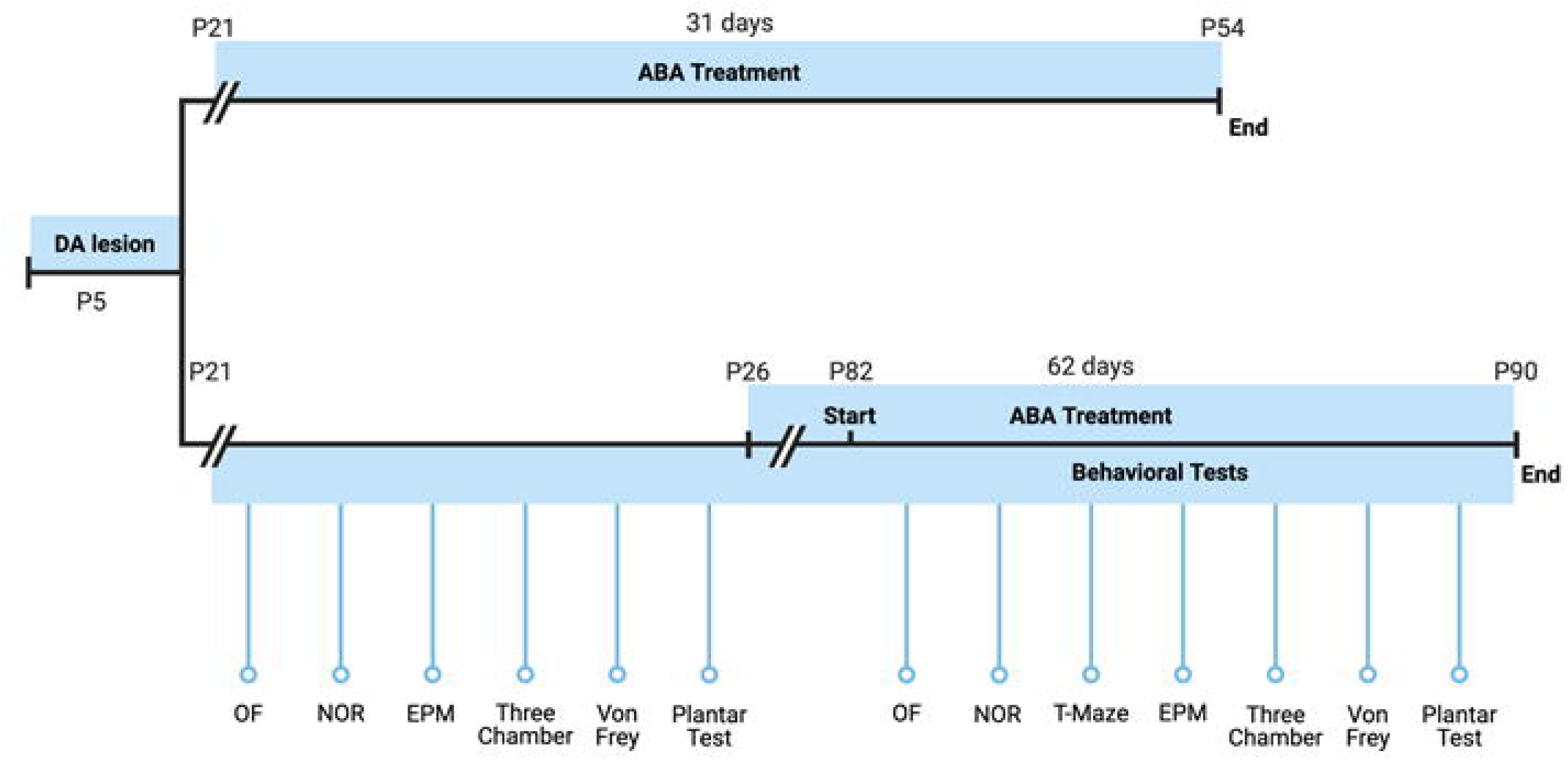
Experiment design timeline. Dopamine lesion (neonatal 6-OHDA injection into lateral ventricle AP −2 mm, ML ±0.6 mm, DV −1.3 mm from Bregma at P5). Baseline started at weaning on P21 until P26. ABA or vehicle administration started on P21 for one month and P26 for two months. Behavioral tests were carried out for one week before terminating the experiment.

### Animals and surgical procedures

Ninety-three Swiss female mice (Janvier-Labs; Saint-Berthevin, France) were kept at the animal facility of the University Jaume I. The procedures followed European Community norms on the protection of animals used for scientific purposes. The experiments were approved by the Ethics Committee of the University Jaume I (scientific procedure 2020/VSC/PEA/0099). The animals were maintained on a 12h:12h light cycle and provided with food and water ad libitum. Pups were housed with their mothers and kept at constant temperature conditions (24°C ± 2). After weaning, animals were housed in groups of 2-4 mice to reduce isolation-induced stress. Ninety-three female mice were randomly assigned to sham, control group of mice undergo surgery with vehicle inoculation, but no lesion; or lesioned groups. At P5, pups received 6-OHDA (Sigma-Aldrich, France) or vehicle in one of the lateral ventricles as described in (45). Thirty minutes before surgery, mice were injected with desipramine hydrochloride pretreatment (20 mg/kg sc; Sigma-Aldrich, France), an inhibitor of the noradrenergic nerves. Mice were anesthetized with 1% isoflurane and maintained with 0.3% isoflurane during surgery. Twenty-five μg of 6-OHDA dissolved in 3 μL ascorbic acid 0.1% or vehicle were infused into one of the lateral ventricles (stereotaxic coordinates: AP −2 mm, ML ± 0.6 mm, DV −1.3 mm from Bregma) (50). After weaning (P21), base line was performed between P21 and P26. After that, from P26 to P90, mice were randomly administered either ABA (20 mg/L) or vehicle (NaOH 0.0008 %) in the drinking water. Animal welfare was monitored all through the procedure, according to Ethical committee rules.

### Behavioral procedures

For all the behavioral paradigms, mice were habituated to the testing room 30 min before performing each test. The tests were conducted during the day, with a dim light. All apparatuses were cleaned using a 30% ethanol solution between trials and animals. Mice performed the paradigms as described previously (46).

*Open Field:* spontaneous locomotor activity and anxiety were assessed through the open field test at P21 and P82. Mice were placed in the open field cage facing one of the walls and allowed to freely explore the arena for 10 min. Distance traveled (cm), speed (cm/s), time in the center (sec) and latency to cross for the first time the center quadrants with all four legs (sec) were quantified. *Novel Object Recognition (NOR):* recognition memory was assessed through NOR test at P22 and P83. Mice were left to explore two identical objects for 10 min (familiarization phase). After 30 min, mice were sent back to the arena and allowed to explore one of the previous objects (familiar) and a novel object for 10 min (test phase). The first 3 minutes were analyzed, and data were expressed as the discrimination index (DI) ((Time exploring the novel object - Time exploring the familiar object)/Total exploration time). In this paradigm, equal or similar exploration of novel and familiar object (d-index = 0) indicates impaired novelty recognition memory.

*T-maze:* spatial memory was assessed through T-maze test at P84. Mice was placed in the starting arm and left to explore for 5 min, with access to two of the three arms (familiarization phase). Mice were then returned to the home cage for a 30 min intertrial interval and then placed back in the starting position but now with free access to all three arms for 5 min (test phase). The arm that was closed during the familiarization phase was considered the “novel” arm, and the arm visited during the familiarization phase was considered the “familiar” arm. Data were expressed as DI ((Entries on the novel arm − Entries on the familiar arm)/Total entries)).

*Elevated Plus Maze (EPM):* impulsivity was assessed through EPM test at P23 and P85. All mice were placed in the center of the maze and allowed to run freely around the maze for 10 min. Data were expressed as the percentage of time spent exploring the open arms compared with the total time in the EPM.

*Three Chamber:* social interaction was assessed through three chamber test at P24 and P86. Animals were placed in the central chamber and allowed to freely explore all chambers for 10 min (habituation phase). Immediately after the habituation phase, we performed the test phase (10 min). Mouse was allowed to explore either an object or a co-specific mice protected by a fence placed in the lateral chambers. The chamber in which the co-specific mice was placed was balanced. In addition, we used different co-specifics mice to balance them. Data were expressed as the percentage of time spent exploring the co-specific mice compared with the total time spent exploring. Climbing or running around it was not considered exploration.

*Von Frey:* nociceptive response to mechanical stimulus was assessed using von Frey setup at P25 and P87. During 30 min, mice were habituated to individual cages with a mesh floor. The plantar surface of the hind paws was stimulated by calibrated von Frey filaments of different grams to set the withdrawal threshold. For both hind paws, three to five measurements were registered, with an interval of 30 s between each. The grams of the filament at which the mouse withdrew its paw was considered to be the mechanical pain threshold value.

### Immunofluorescence procedure

Immunofluorescence was performed as described (51). Briefly, mice were anesthetized and perfused with saline (0.9% NaCl) followed by fixative (4% paraformaldehyde in 0.1M PB, pH 7.4). After perfusion, the brains were removed, postfixed overnight, and cryoprotected in 30% sucrose in 0.01M PBS pH 7.4 for 3 days. The brains were cut in the rostro caudal direction (40μm) using a sliding microtome Leica SM2010R (Leica Microsystems, Heidelberg, Germany). Primary antibodies mouse anti-Tyrosine Hydroxylase (Sigma-Aldrich, France; 1:5000), rabbit anti-Iba1 (FUJIFILM Wako Chemicals Europe GmbH, Deutschland; 1:1000), rabbit anti-vGluT1 (Synaptic Systems, Germany; 1:2000), rabbit anti-vGAT (Synaptic Systems, Germany; 1:1000), mouse anti-Neurofilament-L (Synaptic Systems, Germany; 1:2000), mouse anti-MAP2 (Invitrogen, Waltham, United States; 1:2000), rabbit anti-Homer1 (Synaptic Systems, Germany; 1:2000) and rabbit anti-NLRP3 (Thermo Scientific, Rockford, IL, USA; 1:300) were incubated over-night. Next, sections were rinsed and incubated for 2h at RT with donkey anti-mouse Cy3 or donkey anti-rabbit Alexa 488 secondary antibodies (Jackson Immunoresearch, Suffolk, UK). Finally, sections were mounted on slides and covered using Fluoromount-G mounting medium (Invitrogen, California, USA).

### Imaging and analysis

Fluorescence images were taken with a confocal scan unit with a module TCS SP8 equipped with argon and helio-neon laser beams attached to a Leica DMi8 inverted microscope (Leica Microsystems). Excitation and emission wavelengths for Cy3 were 433 and 560–618nm respectively; Alexa488 labeled excitation wavelength was 488nm and its emission at 510–570nm. For the quantification of Tyrosine Hydroxylase labeling, we used a 10x lens. Image J software was used to count Tyrosine Hydroxylase labeling. Data were expressed as the percentage of Tyrosine Hydroxylase labeling with respect to sham group. For the quantification of Iba1 labeling, we used 20x lens. The custom-designed Image J software macro called “MACROglia” (publicly available at the Github website: https://github.com/SandraSSB/MACROglia_cell-morphology-analysis) combined with FracLac plugin (52) was used to analyze the microglia morphology in sections from sham and 6-OHDA groups of both females and males as previously described (53). The microglia morphological parameters that were analyzed were (I) <u>fractal dimension (D)</u>, this parameter evaluates cellular branching complexity; (II) <u>cell area</u>, meaning the total number of pixels corresponding to the area occupied by the cell, soma, and branches; and (III) <u>cell perimeter</u>, based on the single outline cell shape. For the quantification of double staining vGluT1/NF-L, vGAT/NF-L and Homer1/MAP2, we used 63x lens. Image J software was used to count the number of vGluT1, vGAT and Homer1 points (minimum 3 pixels were considered) in one NF-L or MAP2-positive fiber in 20–25 Z-plane sections from sham and 6-OHDA groups. Ten different axons per animal were analyzed by a researcher blind to the condition. Data is calculated as the number of vGluT1, vGAT and Homer1 positive signals on the NF-L or MAP2 fiber, normalized to the area of the fiber. For the quantification of NLRP3 labeling we used 63x lens. Data were expressed as the mean of gray values per area.

### RNA Extraction and Real-Time Quantitative Polymerase Chain Reaction (RT-qPCR)

Total RNA was extracted from the ACC (n = 51) and homogenized in 350μL of lysis buffer according to the PureLink™ RNA Mini Kit (Thermo Scientific, product no. 12183018A, Rockford, IL, USA). Genomic DNA was removed using a spin-column process during the RNA extraction. In addition, DNAse I treatment (Thermo Scientific, Rockford, IL, USA) was performed to ensure the complete removal of genomic DNA. RNA samples were eluted in 20μL of nuclease-free water and reverse transcribed to cDNA using a High-Capacity cDNA Reverse Transcription Kit (Thermo Scientific, product no. 4368814, Rockford, IL, USA) following the manufacturer’s instructions. RT-qPCR reactions were carried out using Maxima SYBR Green/ROX qPCR MM (Thermo Scientific, product no. K0221, Rockford, IL, USA) in an Applied Biosystems StepOne Plus™ Real-Time PCR System (Foster City, CA, USA). The list of primers is presented in Supplementary Table S1. At the end of each PCR reaction, a melting curve stage was performed to confirm that only one PCR product was amplified in these reactions. The relative gene expression to SEM was calculated by using the 2^-ΔΔCt^ method for each reaction and by using the housekeeping gene GAPDH as internal control.

### Statistical analysis

The analysis was carried out with GraphPad software (GraphPad Prism V9 software, GraphPad, La Jolla, CA, USA). Data were subjected to the Shapiro-Wilk test for Gaussian distribution. If normality was confirmed, data were reported as the mean ± SEM and the "n" the number of independent subjects. Two-tailed paired Student’s t-test with the probability set at α<0.05 was used. Paired t-test was applied to evaluate age and treatment effect in the same individual before and after (P21 vs. P90), and unpaired t-test to evaluate the effect of either lesion or ABA treatment in different animal groups. One-tailed unpaired Student’s t-test with the probability set at α<0.05 was used to analyze the effect of lesion and ABA treatment on the microglia morphology, and cytokines expression of ACC.

## Results

As described previously (46), 6-OHDA injected animals with less than 45% of the VTA area lesioned were discarded from the analysis.

### Neonatal 6-OHDA lesion-induced hyperactivity and impulsivity in young mice continues to adulthood. ABA administration normalizes locomotor and impulsive behavior

Spontaneous locomotor activity was measured by distance travelled and speed during 10 minutes on the open field test. Representative tracking of mice activity is represented (Fig. 2A). We confirmed that 6-OHDA neonatal lesion induces hyperactivity in young female (P21) compared to sham mice [SHAM-VEH P21 vs. 6-OHDA-VEH P21] (Supplementary Table S2). To understand the effect of the lesion and treatment over time; we compared mice behavior at young and adulthood time points as depicted. Lesioned mice continued hyperactive in the adulthood [6-OHDA P21 vs. 6-OHDA-VEH P90] (Fig. 2B; Fig.2C), whereas in control mice, age reduce spontaneous locomotor activity [SHAM P21 vs. SHAM-VEH P90], distance (Fig. 2B; p = 0.0107*) and velocity (Fig. 2C; p = 0.0081**). Indeed, both groups of untreated adult mice [SHAM-VEH P90 vs. 6-OHDA-VEH P90] were significantly different (Supplementary Table S3).

**Figure 2.**
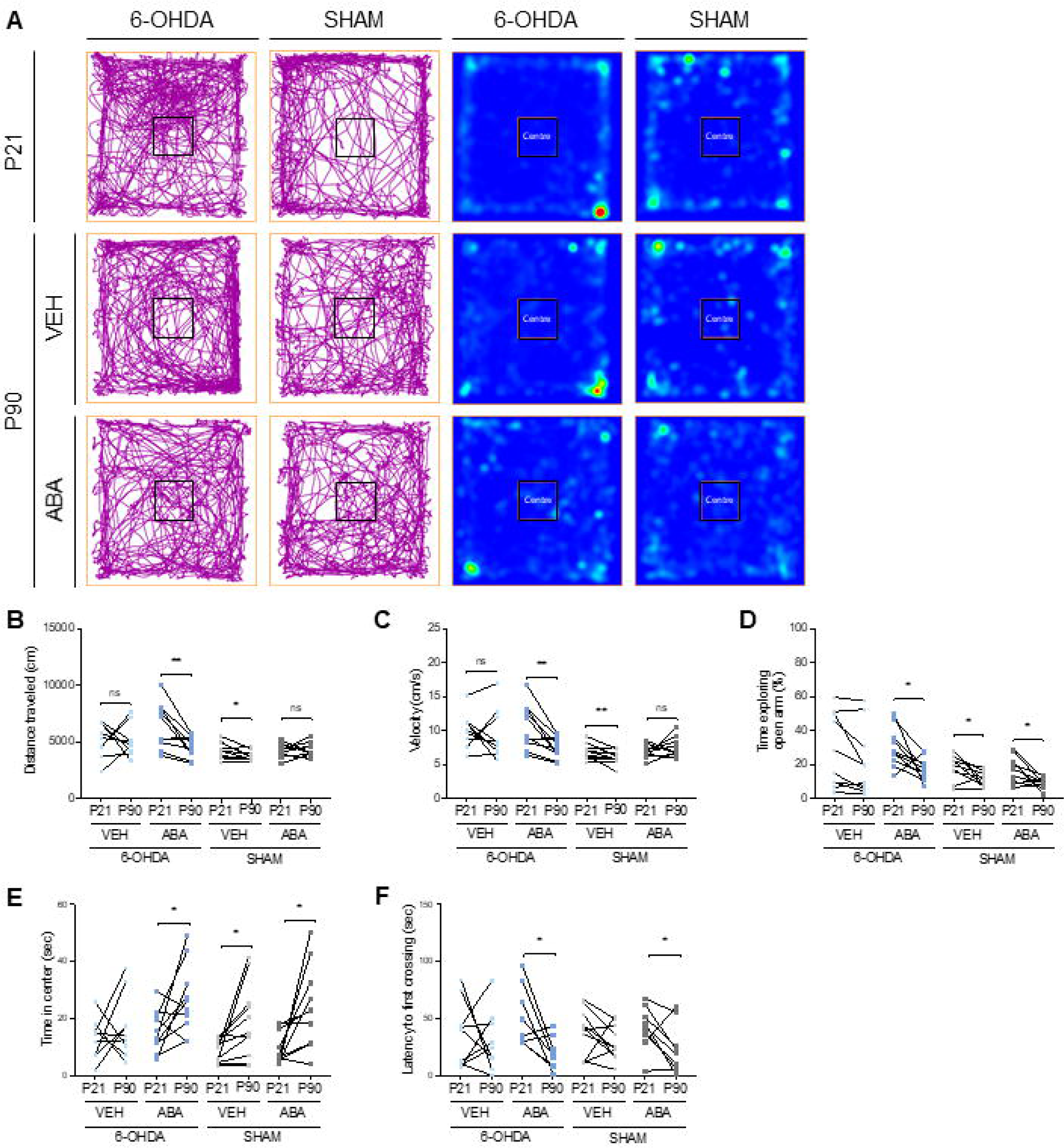
Neonatal 6-OHDA lesion induces hyperactivity, impulsivity in young and adult mice, and anxiety in adults lesioned mice. ABA administration normalizes locomotor and impulsive behavior and alleviates normal development-induced anxiety in adults. **(A)** Representative tracking plot and heat map showing alteration in locomotion and exploratory behavior in open field test. **(B)** Distance travelled (cm) and **(C)** speed (cm/s) in the arena. **(D)** Time exploring open arms in the EPM. **(E)** Time (sec) and **(F)** latency to the first cross into the center (sec) of the arena. Data are presented as mean ± SEM (n = 8-11 per condition) and analyzed by paired two-tailed Student t-test (ns = not significant, * p < 0.05, ** p < 0.01).

ABA administration significantly reduced lesion-induced hyperactivity in adults [6-OHDA P21 vs. 6-OHDA-ABA P90], distance (Fig. 2B; p = 0.0064**) and velocity (Fig. 2C; p = 0.0034**). Interestingly, the age-induced downregulation of spontaneous locomotor activity in control mice was prevented by ABA treatment [SHAM P21 vs. SHAM-ABA P90] in distance (Fig. 2B) and velocity (Fig. 2C). ABA treatment effect was also identified by inter-group comparison [SHAM-VEH P90 vs. SHAM-ABA P90] (Supplementary Table S4).

These findings suggest that ABA treatment can contribute to regulate locomotor activity, by reducing lesion-induced hyperactivity, and preventing the reduction in spontaneous movement, associated with physiological development.

We next analyzed the effects caused by the neonatal dopaminergic lesion on impulsivity by evaluating behavior at the EPM performance. This paradigm is validated to measure impulsivity in ADHD (54) and in aging mice models (55).

In young mice, neonatal 6-OHDA lesion increased significantly the time exploring the open arms compared to control mice [SHAM P21 vs. 6-OHDA P21] (Supplementary Table S2). This impulsive behavior continued to adulthood [6-OHDA P21 vs. 6-OHDA-VEH P90] (time exploring the open arms, Fig. 2D), and ABA treatment reduced impulsive behavior in neonatally lesioned mice [6-OHDA P21 vs. 6-OHDA-ABA P90] (time exploring the open arms, Fig. 2D; p = 0.0178*). Normally developed females further reduced impulsivity with age [SHAM P21 vs. SHAM-VEH P90], as illustrate by reduced time exploring (Fig. 2D; p = 0.0178*).

These results indicate that impulsivity, measured as the time spent in open arms, is higher at young ages compared to adults, and it is greatly exacerbated by the neonatal lesion. ABA can effectively modulate impulsive behavior, by reducing it in lesioned mice, and preventing age-associated locomotor activity decline in controls.

### ABA treatment alleviates neonatal 6-OHDA lesion-induced anxiety in adults

Anxiety was measured in the open field test by the time and latency to first crossing into the center area, parameters that are associated to anxious behavior. Representative tracking and heat map are depicted in Fig. 2A. In young mice, neonatal 6-OHDA lesion increased time in the center, and reduced the latency to first crossing, [SHAM P21 vs. 6 OHDA P21], (Supplementary Table S2). This observation may be interpreted as reduced anxiety, but this behavior could be due to the hyperactivity and impulsivity, characteristics that are also associated to dopaminergic lesion.

Normal developed mice [SHAM P21 vs. SHAM-VEH P90], increase time in center with age (Fig. 2E; p = 0.0199*) (whereas decreasing locomotor activity Fig. 2B-C), indicating reduced anxious behavior. This finding supports the notion that normal development can contribute to decrease anxiety. Neonatal dopaminergic lesion impairs the age effect observed in control mice [6-OHDA P21 vs. 6-OHDA-VEH P90].

ABA treatment in lesioned females [6-OHDA P21 vs. 6-OHDA-ABA P90] rescues it, significantly increasing the time in the center (Fig. 2E; p = 0.0390*) and decreased the latency to first crossing into the center (Fig. 2F; p = 0.0238*). Whereas in ABA treated controls did not differ much from vehicle treated controls, as far as decreased time in the center [SHAM P21 vs. SHAM-ABA P90] (Fig. 2E; p = 0.0204*) and decreased the latency to first crossing (Fig. 2F; p = 0.0386*).

These results strongly support that normal development decreases anxious behavior, but not in neonatal dopamine lesion mice. ABA treatment of lesioned mice rescues it, therefore providing a potential effective anxiolytic tool for ADHD patients.

### Mechanical sensitivity is increased by neonatal dopaminergic lesion. Age and ABA treatment alleviates mechanical hypersensitivity in lesioned mice

Mechanical sensitivity threshold was measured in the Von Frey test. Young lesioned females were hypersensitive compared to control mice [SHAM P21 vs. 6-OHDA P21] (Supplementary Table S2, p < 0.0001****). Development increased the threshold in lesioned mice [6-OHDA P21 vs. 6-OHDA-VEH P90] (Fig. 3A; p = 0.0405*); whereas decreased it in control mice [SHAM P21 vs. SHAM-VEH P90] (Fig. 3A; p = 0.0025**), suggesting that development may intensify the sensitivity to mechanical stimulus in control subjects, but eases the hypersensitivity in ADHD subjects. Two-months of ABA treatment augmented the threshold in lesioned mice [6-OHDA P21 vs. 6-OHDA-ABA P90] (Fig. 3A; p < 0.0001****), indicating that ABA further alleviates hypersensitivity in ADHD adults. Indeed, there is a significant difference between [6-OHDA-VEH P90 vs 6-OHDA-ABA P90] (Supplementary Table S5, p < 0.001 ***). Very interestingly, ABA in control mice prevented the reduction in threshold [SHAM P21 vs. SHAM-ABA P90] (Fig. 3A), indicating that ABA may contribute to reduced sensitivity in adult.

**Figure 3.**
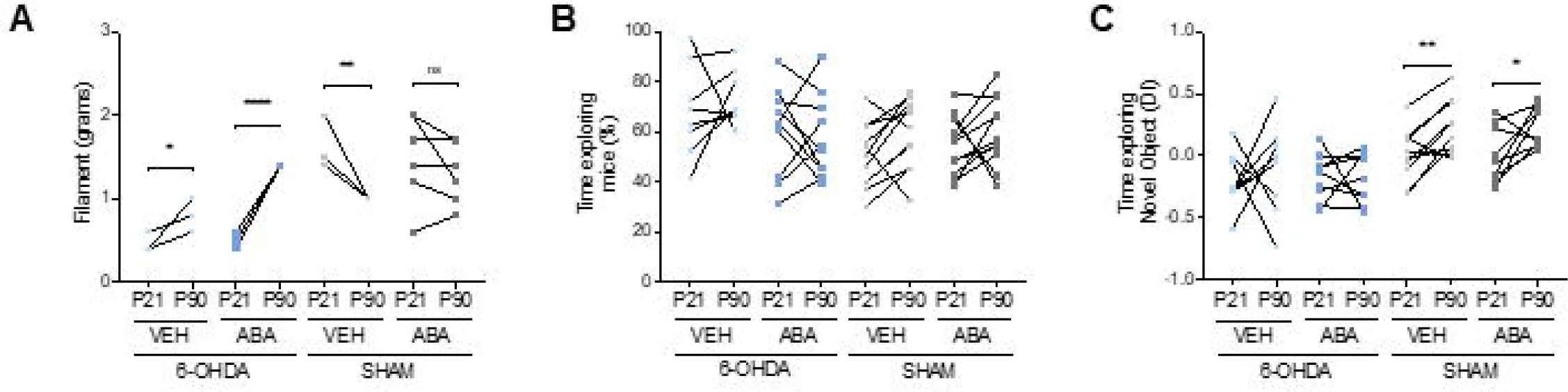
Neonatal 6-OHDA lesion increased mechanical sensitivity and social interaction and impairs recognition memory. Age and ABA treatment alleviates mechanical hypersensitivity in lesioned mice. **(A)** Mechanical threshold (grams of filament) in Von Frey test. **(B)** Time exploring the co-specific mice expressed as the percentage of total time exploring. **(C)** Time exploring (d-index) the novel object in the novel object recognition test. Data are expressed as a DI ((Time exploring novel – time exploring familiar)/total time exploring), presented as mean ± SEM (n = 4-11 per condition) and analyzed by paired two-tailed Student t-test (ns = not significant, * p < 0.05, ** p < 0.01, **** p < 0.0001).

These findings indicate that ABA treatment effectively downregulate mechanical hypersensitivity in ADHD subjects, but also in normally developed mice.

### Neonatal 6-OHDA lesion increased social interaction. ABA administration normalizes this effect

Social interaction (SI) was measured by co-specific mice exploration. We found no changes in SI during development in any condition (Fig. 3B). However, when the 6-OHDA lesion effect was analyzed between groups, we observed that lesion in young females 6-OHDA lesion renders mice with higher social activity [SHAM P21 vs. 6-OHDA P21] (Supplementary Table S2; p < 0.01**). An effect that was maintained through adulthood [SHAM-VEH P90 vs. 6-OHDA-VEH P90] (Supplementary Table S3; p < 0.05*). These results showed that neonatal 6-OHDA lesion may increase sociability in young and adults. Curiously, in lesioned adults, ABA treatment significantly reduced the hyper-sociability [6-OHDA-VEH P90 vs. 6-OHDA-ABA P90] (Supplementary Table S5; p < 0.05*). These results suggest that ABA treatment counteracts the DA deficit effect in sociability.

### Neonatal 6-OHDA lesion impairs recognition and spatial memory in adult mice. ABA does not rescue this effect

Next, we aim to understand if neonatal 6-OHDA lesion affects novel recognition memory in young and adult female mice through NOR test. None of the infant mice learned (d-index < 0.2), but whereas normally developed mice significantly improved novel recognition performance with age [SHAM P21 vs. SHAM-VEH P90] (Fig. 3C, p = 0.0057**), lesioned females did not [6-OHDA P21 vs. 6-OHDA-VEH P90] (Fig. 3C). Moreover, ABA did not rescue lesion-induced impairment [6-OHDA P21 vs. 6-OHDA-ABA P90] (Fig. 3C). Comparing lesion effect amongst groups, further confirmed the deleterious effect of neonatal 6-OHDA lesion in adult mice [SHAM-VEH P90 vs. 6-OHDA-VEH P90] (Supplementary Table S3; p < 0.05*).

These results showed that normal development improves performance in the novel recognition task, but it is impaired by neonatal dopaminergic lesioned mice, and it cannot be rescued by ABA treatment.

Furthermore, spatial memory was analyzed in adult mice through T-Maze (Supplementary Fig. S1A). Young, P21 mice were not analyzed in this test, since they are too young to perform optimally (as we confirmed in the NOR test). We observed that neonatal lesion results in adults with reduced number of entries in the novel arm compared to sham females [SHAM-VEH P90 vs. 6-OHDA-VEH P90] (Supplementary Fig. S1B; p < 0.01**). Similarly, ABA treatment did not recover the effect on 6-OHDA lesion [6-OHDA-VEH P90 vs. 6-OHDA-ABA P90] (Supplementary Fig. S1B), thus, DA deficit impairs T-maze performance, suggestive of impaired spatial memory or lack of novelty recognition, but ABA treatment cannot rescue it.

### Neonatal 6-OHDA lesion promotes microglia polarization to proinflammatory status in pIC, ACC and hippocampus. Two-months of ABA treatment restores microglia morphology

Microglia of three-month old mice (Fig. 4A) were visualized in several brain areas: the pIC, ACC, and dentate gyrus of hippocampus (Fig. 4B) using Iba-1 staining (Fig. 4C). Morphological changes of microglia in these areas were evaluated by fractal, area, and perimeter parameters. Neonatal 6-OHDA lesion [SHAM-VEH P90 vs. 6-OHDA-VEH P90] significantly reduces the complexity of the microglia branching in the <u>insular cortex</u> in 3-month-old female mice, reducing the microglia fractal (Fig. 4D; p = 0.0001***), area (Fig. 4E; p = 0.0008***), and perimeter (Fig. 4F; p = 0.0093**). Similar effect was observed in <u>ACC</u> microglia, fractal (Fig. 4G; p = 0.0006***), area (Fig. 4H; p = 0.0053**), and perimeter (Fig. 4I; p = 0.0095**); and <u>hippocampus</u> microglia, fractal (Fig. 4J; p = 0.0017**), area (Fig. 4K; p < 0.0001****) and perimeter (Fig. 4L; p = 0.0003***).

**Figure 4.**
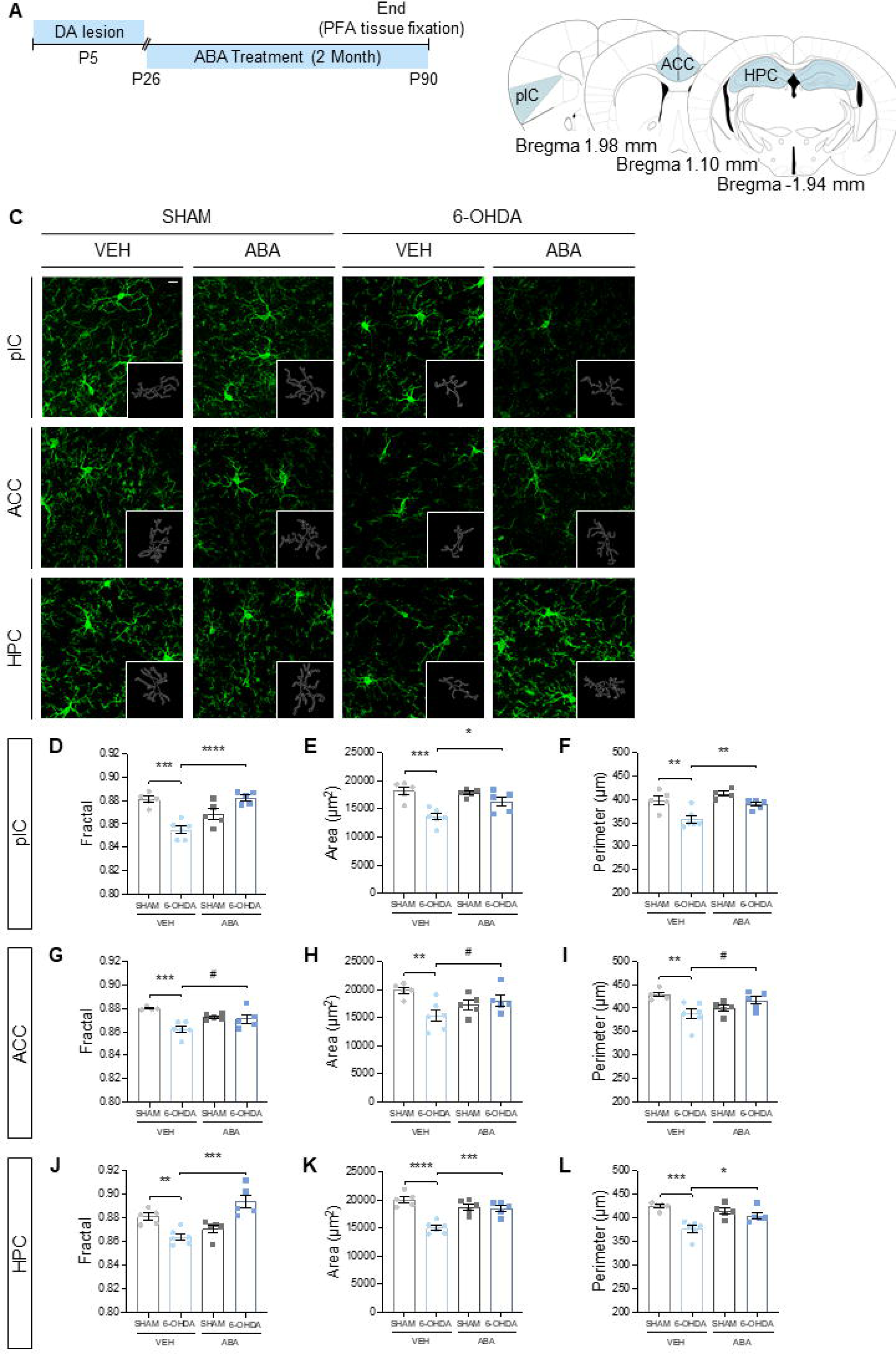
Dopaminergic lesion activates microglia and ABA treatment prevents activation. **(A)** Schematic experimental design for dopaminergic lesion (6-OHDA) indicating the extension of ABA treatment and the moment of brain sample collection to evaluate the neuroinflammation through microglia morphology. **(B)** Schematic representation of the different area that were studied. **(C)** Representative confocal microscopy images from pIC, ACC, and Hippocampus showing Iba1 marker. Inserts in every image are an outline of the microglia cell in the corresponding image. Calibration bar; 10 µm. **(D)** Fractal **(E)** area (µm^2^) and **(F)** perimeter (µm) in pIC. **(G)** Fractal **(H)** area (µm^2^) and **(I)** perimeter (µm) in ACC. **(J)** Fractal **(K)** area (µm^2^) and **(L)** perimeter (µm) in Hippocampus. Data are expressed as mean ± SEM (n = 4-6 per condition, with 5 cells analyzed per animal) and analyzed by unpaired two-tailed Student t-test (* p < 0.05, ** p < 0.01, *** p < 0.001, **** p < 0.0001) and unpaired one-tailed Student t-test (#p < 0.05).

We confirmed that 2-month of ABA treatment rescued neonatal lesion effect [6-OHDA-VEH P90 vs. 6-OHDA-ABA P90] in the <u>insular cortex</u> increasing fractal (Fig. 4D; p < 0.0001****), area (Fig. 4E; p = 0.0232*) and perimeter (Fig. 4F; p = 0.0098**), also in the <u>ACC</u> fractal (Fig. 4G; p = 0.046#), area (Fig. 4H; p = 0.0482#) and perimeter (Fig. 4I; p = 0.0346#); and in <u>hippocampus</u> fractal (Fig. 4J; p = 0.0006***), area (Fig. 4K; p = 0.0006***) and perimeter (Fig. 4L; p = 0.0230*).

Neonatal dopaminergic lesion affects microglia morphology in diverse brain areas (46) of young mice (two-month-old). Presently, data confirms that this alteration lasts until adulthood (three-month-old). ABA, short and longer treatment (one and two-month) leads to higher ramified morphology, suggesting a control healthy microglia stage.

### Dopaminergic lesion increase M1 cytokines and decrease M2 marker in young mice and ABA normalizes the effect

To further understand the mechanism by which dopaminergic lesion and ABA exerts the behavioral effects, molecular alterations in ACC were analyzed at one and two-months of ABA treatment (Fig. 5A).

**Figure 5.**
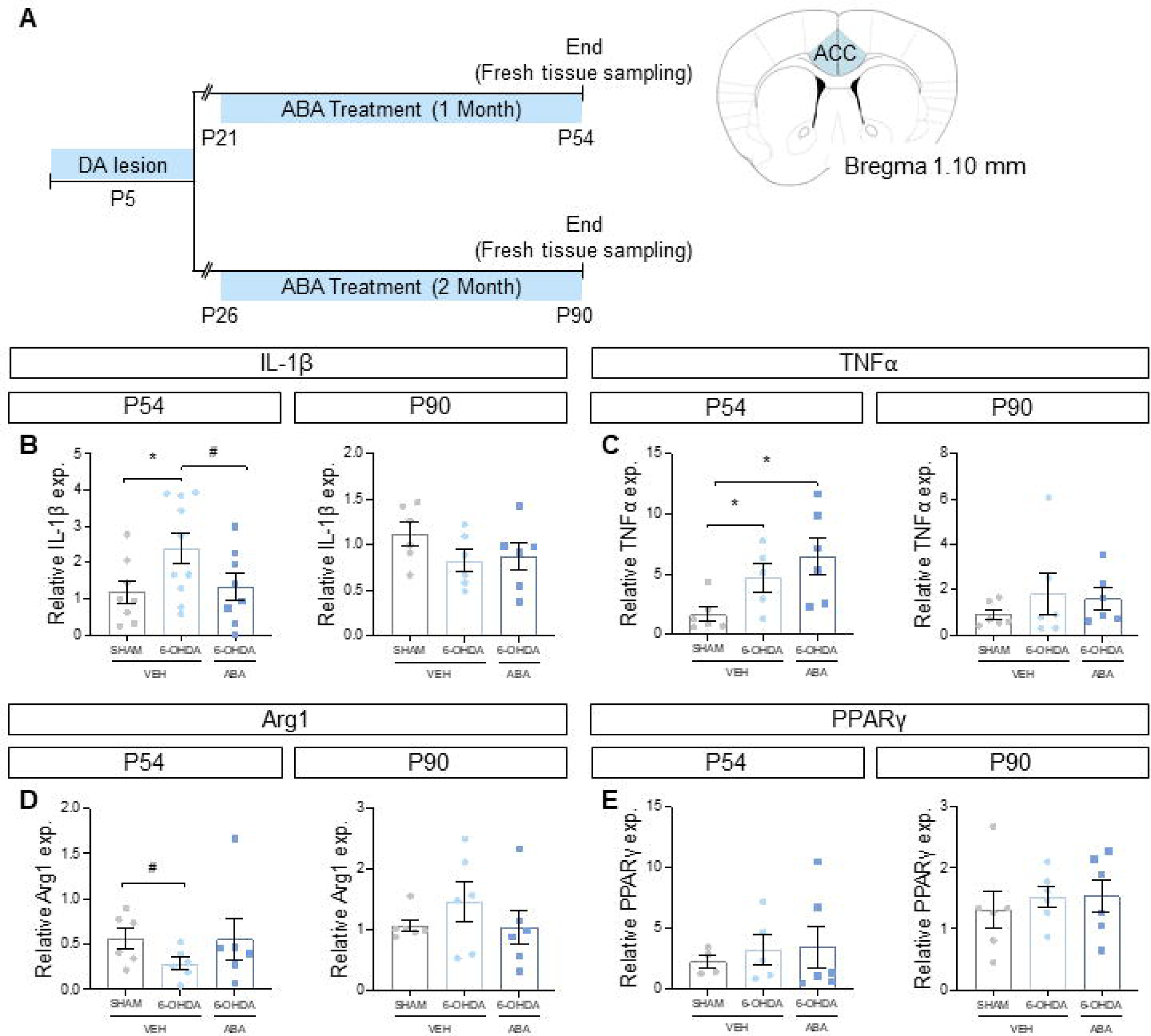
Neonatal 6-OHDA lesion increase pro-inflammatory cytokines expression and decrease anti-inflammatory Arg1 expression. **(A)** Schematic experimental design for dopaminergic lesion (6-OHDA) indicating the extension of ABA treatment and the moment of brain sample collection to evaluate pro-inflammatory and ant-inflammatory cytokines expression. Schematic representation of the area that was studied. **(B)** IL-1β, **(C)** TNFα, **(D)** Arg1 and **(E)** PPARγ expression at P54 (1 month of ABA treatment) and P90 (2 months of ABA treatment). Data are presented as mean ± SEM (n = 4-10 per condition) and analyzed using two-tailed Student t-test (* p < 0.05) and one-tailed Student t-test (# p < 0.05).

Pro-inflammatory cytokines IL-1β and TNFα expression increased in young animals that has undergone 6-OHDA lesion [SHAM-VEH P54 vs. 6-OHDA-VEH P54], IL-1β (Fig. 5B; p = 0.0437*) and TNFα (Fig. 5C; p = 0.0347*). One-month of ABA treatment [6-OHDA-VEH P54 vs. 6-OHDA-ABA P54], reduced significantly the expression of IL-1β (Fig. 5B; p = 0.0422#), but not TNFα (Fig. 6D). Curiously, these differences were not maintained in adults [SHAM-VEH P90 vs. 6-OHDA-VEH P90], in IL-1β (Fig. 5B) and TNFα expression (Fig. 5C).

**Figure 6.**
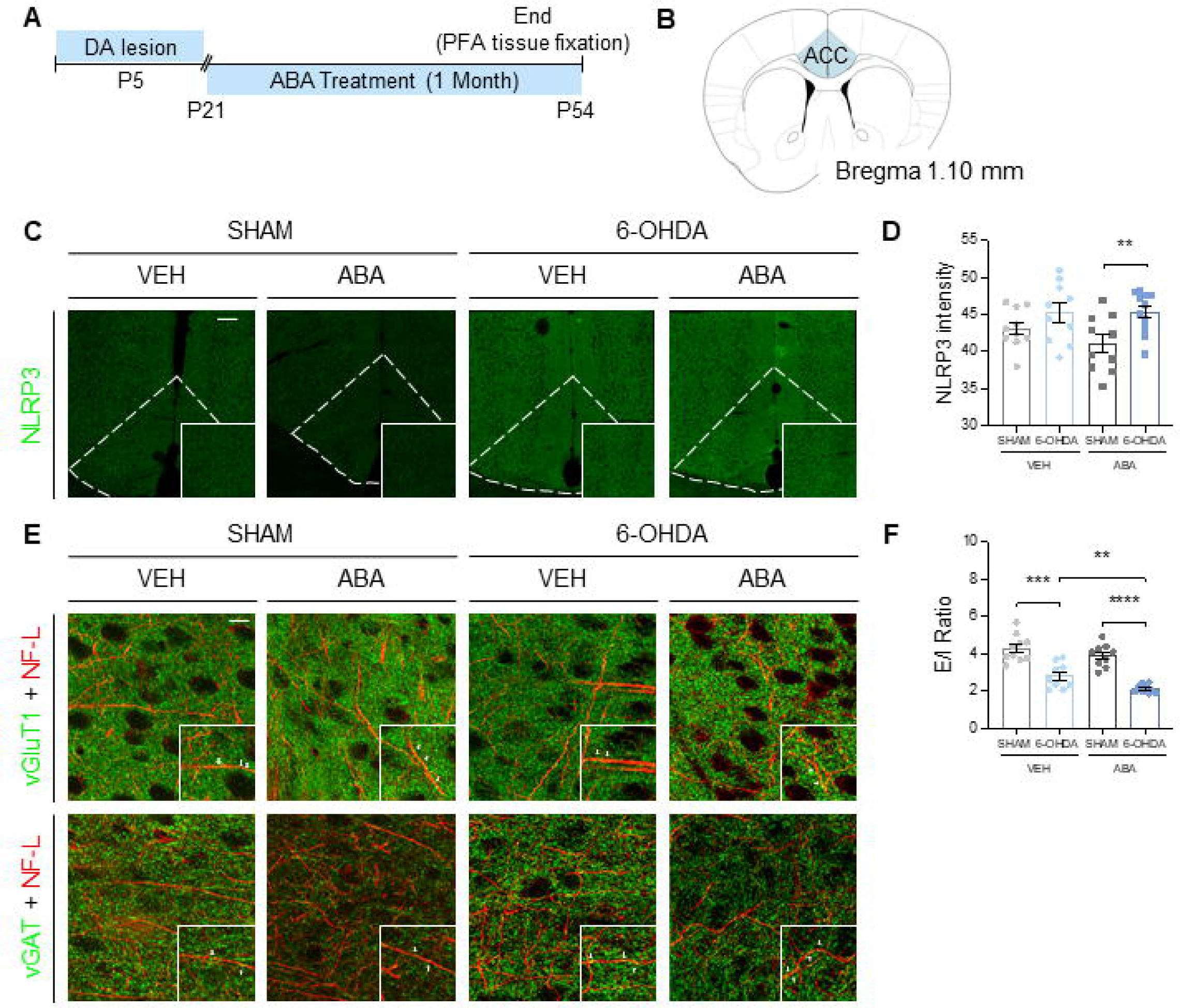
Dopaminergic lesion impairs NLRP3 and E/I ratio. **(A)** Schematic experimental design for dopaminergic lesion (6-OHDA) indicating the extension of ABA treatment and the moment of brain sample collection to evaluate the neuroinflammation and E/I ratio. **(B)** Schematic representation of the area that was studied. **(C)** Representative confocal microscopy images from ACC showing NLRP3 marker. Calibration bar; 100 µm. **(D)** NLRP3 intensity (mean gray value). **(E)** Representative confocal microscopy images from ACC showing vGluT1, vGAT and NF-L markers. Inserts in every image show vGluT1 and vGAT in pre-synaptic terminal. Calibration bar; 100 µm. **(F)** E/I ration in ACC females. Data are presented as mean ± SEM (n = 9-11 per condition) and analyzed using two-tailed Student t-test (** p < 0.01, *** p < 0.001, **** p < 0.0001).

**Figure 7.**
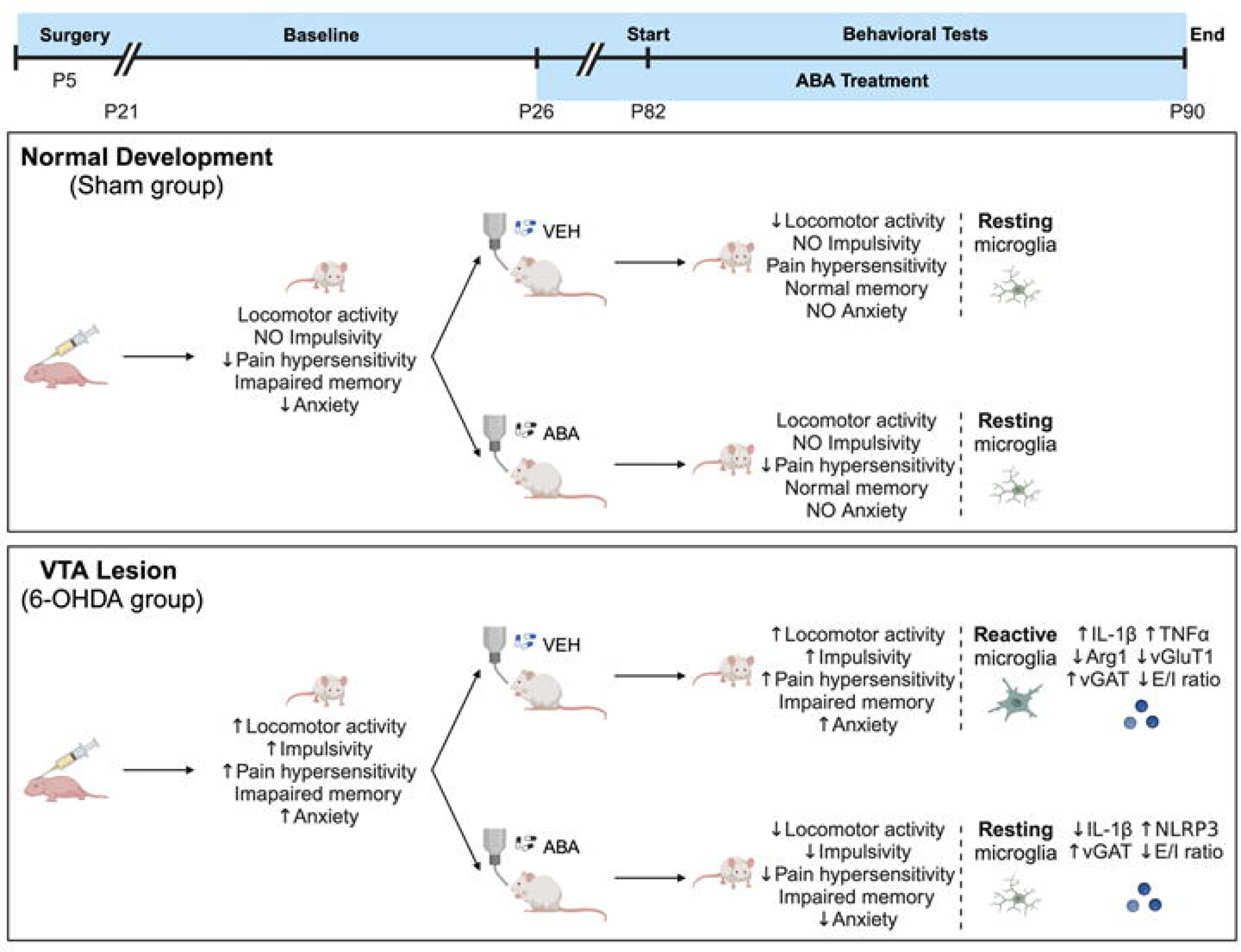
Graphical Abstract. Effect in adult females of neonatal dopamine depletion and ABA treatment. Neonatal 6-OHDA dopaminergic lesion of the VTA induces behavioral changes hyperactivity, impulsivity, hypersensitivity and increased social interaction in P21 and P90 females, and memory impairment in P90. Two-months of ABA treatment improved hyperactivity, impulsivity, anxiety, hypersensitivity, and alterations in social interaction, but not cognitive impairment. In the ACC of young adult mice (P60) dopamine deficiency induced mRNA alteration (as indicated); and E/I imbalance. ABA treatment restored microglia morphology, IL-1β expression, and increased vGAT levels. Black arrows indicate changes induced by age or treatment, and blue arrows indicate changes induced by lesion at P21, * indicate that anxious behavior is masked by hyperactivity and impulsivity.

On the other hand, the expression of Arg1, a marker of M2 microglia, was reduced in young, lesioned females [SHAM-VEH P54 vs. 6-OHDA-VEH P54] (Fig. 5D; p = 0.0296#), but not in adults (Fig. 5D). Interestingly, one-month of ABA treatment abrogates this difference, suggesting that ABA prevents Arg1 reduction in lesioned mice.

Finally, the mechanism by which ABA acts was investigated. ABA have been shown to activate PPARγ expression in vivo (56), but we found not expression alterations with ABA [6-OHDA-VEH P54/P90 vs. 6-OHDA-ABA P54/P90] (Fig. 5E).

These results suggest that in young mice microglia displays a clear pro-inflammatory status (M1), with increased expression of IL-1β and TNFα and a reduction of Arg1 (M2). This early inflammatory status may determine long-lasting behavioral alterations.

### NLRP3 inflammasome expression is not significantly altered by 6-OHDA lesion

To further determine these early alterations observed in cytokines and given the relationship between dopamine and NLRP3 inflammasome (57) and the physiopathology of NLRP3 inflammasome overactivation (58,59); we measured NLRP3 expression in young female mice ACC (Fig. 6A-B) by immunofluorescence staining (Fig. 6C). Quantification of fluorescence intensity showed no significant changes in NLRP3 expression in untreated lesioned females [SHAM-VEH P54 vs. 6-OHDA-VEH P54] (Fig. 6D). Lesion effect was only evidenced in ABA treated animals, since the ABA-treated sham displayed lower NLRP3 intensity [SHAM-ABA P54 vs. 6-OHDA-ABA P54] (Fig. 6D; p = 0.0075**).

### Neonatal 6-OHDA lesion alters the excitatory/inhibitory ratio. One month of ABA treatment increased vGAT. At postsynaptic level, no alterations are observed in any condition

Next, based on the important role of excitation/inhibition (E/I) ratio imbalance in neurodevelopmental disorders (60) and the strict control of microglia on E/I balance (61), we hypothesized that in our model microglia alterations could alter E/I balance, and this could be rescued by ABA. To that end we analyzed vesicular glutamate marker (vGluT1) and vesicular GABA transporter (vGAT), vesicular markers that correlate with E/I ratio (62). vGluT1 and vGAT immunodetection were colocalized with the axonal marker Neurofilament-L (Fig. 6E).

We observed that neonatal 6-OHDA lesion reduced significantly the E/I ratio [SHAM-VEH P54 vs. 6-OHDA-VEH P54] (Fig. 6F; p = 0.0003***), due to reduced vGluT1 (Supplementary Fig. S2A; p < 0.05*) and increased vGAT (Supplementary Fig. S2B; p < 0.05*). Remarkably, one-month of ABA treatment further reduced the E/I ratio [6-OHDA-VEH P54 vs. 6-OHDA-ABA P54] (Fig. 6F; p = 0.0076**), significantly increasing vGAT (Supplementary Fig. S2B; p < 0.001***), not affecting vGluT1 (Supplementary Fig. S2A).

We further investigated changes at **postsynaptic** level, evaluating the colocalization of the protein Homer1, which regulates the function of metabotropic glutamate receptor (63) and MAP2, associated with microtubules and enriched in dendrites (Supplementary Fig. S2C). No alterations with 6-OHDA, nor ABA administration were observed (Supplementary Fig. S2D).

These findings suggest that neonatal 6-OHDA lesion alter E/I balance. ABA administration increased the inhibitory vGAT, correlating with the rescue of the microglia morphology providing a potential mechanism of ADHD treatment.

## Discussion

Female neonatal dopaminergic lesion mice model of ADHD remains significantly underrepresented in preclinical studies; thus, we used female mice to analyze the impact of age and anti-inflammatory treatment on symptomatology by comparing behaviors at P21 with P90. As per Semple et al., these time points are equivalent to human brain maturity at approximately 3-year-old and over 20 years old, respectively (49). In this study, we demonstrate that neonatal dopaminergic lesion induces long-term effects. These effects either manifest at younger time points mice and persist throughout adulthood (hyperactivity, impulsivity, and hypersensitivity to mechanical stimuli), or they emerge in adulthood (anxiety and cognitive impairment). Moreover, we demonstrate that ABA treatment alleviates symptoms, except cognitive impairment. We have previously shown the potential anti-inflammatory effects benefits of ABA in different animal models, metabolic syndrome (64) and Alzheimer (53). Our study (46) and others, strongly suggest a link between neuroinflammation with hyperactive or impulsive behavior in ADHD models (65). Therefore, ABA was used as a potential therapy as chronic treatment.

Dopaminergic lesion induced hyperactivity and impulsive behavior in young female mice, compared to controls, similar to the observations reported in other ADHD models utilizing male mice (66,67). Controls, on the other hand, upon reaching adulthood experienced a reduction in spontaneous locomotor activity and impulsivity, aligning with human behavior as prefrontal cortical structures mature (68). With age, in lesioned females, hyperactivity and impulsivity persist, and this persistence echoes the behavior of some ADHD patients, where impulsivity remains a challenge through adulthood, potentially elevating the risk of engaging in activities such as substance abuse disorders or automobile accidents (69). Two-months of ABA treatment effectively reduced hyperactivity and impulsivity in lesioned female mice, consistent with our previous results (46), and with other studies in locomotor alterations models, such as, Parkinson’s disease (70) and in harmaline-induced motor disabilities (71) that have shown the beneficial ABA effect rescuing motor alterations. Hyperactivity and impulsivity have been reported to be regulated by dopamine deficiency and/or insufficient activation of the dopamine D1 receptor in the ACC (72–74). ABA treatment did not alter VTA dopaminergic signal (46), thus arguing for the potential therapeutic strategy targeting downstream mechanism elicited by DA deficiency underlying ADHD symptoms. Remarkably, ABA counteracted age effect in controls, maintained young mice activity, which may be suggestive of a revitalizing effect of ABA. To our knowledge this is the first report of potential rejuvenating ABA effect on healthy subjects.

Anxiety in rodents has been traditionally evaluating their behavior in the open field paradigm. Increased time in center of the arena and reduced latency to the first crossing is associated to reduced anxiety. Whether dopaminergic lesions affect anxiety can be difficult to assess with this paradigm, since the time in center and latency can also be elicited by hyperactivity and impulsivity. Our experimental design allowed us to discern between these possibilities. With age, control mice increased time spent in the center along with decreased locomotor activity and impulsivity, this would indicate less anxiety in adults. Meanwhile, lesioned mice maintain hyperactivity, but also spent more time in the center and reduced latency into the first crossing. When lesioned mice were treated with ABA they increased time in center and reduced latency, while reducing hyperactivity and impulsivity. Taking all these observations together, we postulate that young, lesioned mice spend more time in the center and less time into the first crossing, maybe due to their hyperactive and impulsive tendencies, rather than reduced anxiety. Meanwhile, in adult lesioned mice, ABA treatment reduces anxiety. Our data aligns with previous studies where ABA exerts anxiolytic effect via Protein kinase C (75), ERK signaling (76). This may be explained by the fact that neuroinflammation contributes to anxious and depressive behavior (77). In particular, neuroinflammation in ACC is associated with mood disorders (78,79), also activity in posterior insular cortex correlates with heart-induced anxiety (80,81). Furthermore, our results are very relevant for ADHD management in patients since anxiety occurs as a comorbid symptom in approximately 50% of cases studied (82), providing an alternative therapeutic tool.

Another symptom co-occurring with ADHD is the hypersensitivity to different stimulus, found more often in girls (83) and women (84) patients. Indeed, neonatal dopaminergic lesion induces mechanical hypersensitivity in young females compared to controls, consistent with previous findings (46,85). Interestingly, in controls sensitivity to mechanical stimuli increased with normal development, aligning with findings reported in healthy humans (86), where threshold decreases with age. Conversely, age mitigated hypersensitivity in lesioned mice. Remarkably, ABA treatment decreased sensitivity to mechanical stimuli in both lesioned and control mice, counteracting both lesion effect and age effect in controls. This data aligns with recent studies demonstrating ABA’s ability to alleviate neuropathic pain by downregulating inflammation in the spinal cord (87). Also, neuroinflammation in brain areas is related to pain (88), thus, we hypothesize that ABA’s reduction of neuroinflammation within the ACC circuitry can effectively counteract hypersensitivity.

Hyper-sociability has been noted in males of the spontaneous hypertensive model of ADHD (89). Similarly, in this study sociability was found increased in young, lesioned females, a trait that persisted into adulthood. Although social difficulties are reported for ADHD patients (90), the equivalence to human behavior is a complex issue, and further studies are needed to comprehend the connection between dopamine deficits and the dysregulation of striatal neuropeptides associated with sociability, such as, arginine, vasopressin and oxytocin; which are essential for social behavior (91,92). Another possible explanation is that impulsivity and hyperactivity increased the time exploring the co-specific mice. Consistently, ABA treatment counteracted the hyper-sociability in lesioned female’s lesion, as it regulates locomotor activity.

Cognitive function may be compromised in ADHD patients (93,94). Our study supported that DA deficit impairs spatial working memory using two paradigms, NOR and T-maze. However, ABA did not recover impaired memory, in contrast with our previous reports in different animal models (53,64). This result may be because these paradigms use novelty as the stimulus to learn. Dopamine is required to novelty recognition (95,96), thus we argue that if ABA does not alter dopamine levels, this would indicate different etiology of symptoms. Further research is warranted to fully decipher the potential therapeutic role of ABA in attentional processes, that do not require hippocampal VTA circuitry for novelty.

To understand the mechanism by which ABA exerts these effects we evaluated microglia morphology in the ACC in adult animals. Microglia morphology reflects its activation status (97–99), and we confirmed that neonatal 6-OHDA lesion induced microglia activation in adult animals, demonstrating a potential long-lasting effect on microglia. ABA treatment effectively rescued the microglia morphology in all areas studied, in line with our previous observations (46,53). Moreover, in this study ACC microglia morphology correlated with behavior. However, in memory related tests, microglia morphology of hippocampus appear ramified but not cognitive improvement respect to untreated mice were observed.

Evaluating the microglia status, we found that in young adult (two-months of age) pro-inflammatory IL-1β and TNFα mRNA expression was increased while Arg1, marker of M2 was decreased by neonatal dopamine lesion. Remarkably, ABA treatment recued IL-1β and Arg1, but not TNFα mRNA expression. In older mice (P90) these cytokines alterations were not found, despite the behavioral alterations reported. These results support the role of early microglia alterations inducing long lasting behavioral alterations, as it is reported for other developmental disorders (100). Our data supports that early intervention targeting microglia with ABA is a potential therapeutic alternative with great potential to alleviate adult behavior.

To further decipher ABA mechanism of action we measured PPARγ mRNA expression in young adult female brain. ABA action is mediated by direct binding to the Lanthionine synthetase component C-like protein 2 (LANCL-2) (101,102), that can activate PPARγ function (103). Both mechanisms are strongly linked to inflammatory regulation (56). We found that ABA did not increases PPARγ expression, in contrast to the early reports in spleen (56). This discrepancy may be due to time window of action, or the different tissue examined. These findings cannot rule out activation of PPARγ.

IL-1β protein maturation to active form is mediated by NLRP3 inflammasome (104). DA deficit has been shown to activate NLRP3 in primary human microglia and in a mouse model of Parkinson’s disease (105). We hypothesized that NLRP3 expression in the ACC would be increased by dopamine deficit, aligning with microglia morphology alterations. However, we did not observe that change. Curiously, ABA reduced NLRP3 expression in control but not lesioned mice. This result indicates that DA deficit does not increase NLRP3 expression in the ACC. However, the activity of NLRP3 cannot be specifically determined since the PCR detects IL-1β mRNA expression, not active protein.

Beyond the inflammatory properties, microglia activity is key in the regulation of the excitation/inhibition (E/I) ratio (106), and the E/I imbalance has been found as a fundamental factor underlying a variety of neuropsychiatric and neurodevelopmental disorders (107). In fact, E/I imbalance in ACC and insula (108) is associated to higher nociception animal models of neurodevelopmental disorders (109). In our model, we observed an E/I ratio imbalance, however, contrary to other studies with this animal model, where an increase in hyper-excitability is observed (85), we found a reduced vGluT1 and an increase vGATs, that would suggest hypo-excitability. Interestingly, pro-inflammatory microglia increase glutamate release (110–112) under inflammatory conditions, thereby causing excitotoxicity (regardless of the presynaptic neuron vGluT1 expression). In addition, dopaminergic dysfunction can impair EAATs channels activity in astrocytes and neurons, decreasing the glutamate reuptake from the synapse (113).

Remarkably, we found that ABA treatment increases vGAT, without altering vGluT1. This finding agrees with studies showing that activating the GABAergic tone in the ACC of a rat model of chronic inflammatory pain, alleviated pain and pain-induced anxiety (114,115). Other reports have shown that ABA can facilitate GABAA function, via PPARγ activation (116). Further studies with calcium imaging and electrophysiology would be required to ascertain the role of ABA in neuron excitability.

In summary, our results suggest a dopamine deficit-induced neuroinflammation (105,117), as the triggering factor of hyperactivity, impulsivity, as well as a hypersensitivity. ABA, a well-known anti-inflammatory agent (46), alleviated these symptoms, concomitantly with improved microglia morphology and rescuing cytokines levels indicative of reduced M1 (IL-1β) and increased M2 phenotype (Arg1) at early stages. Moreover, ABA may influence the E/I ratio by increasing vGAT, likely as a result of regulating microglia activity.

### Conclusions

ADHD has been traditionally associated with dopaminergic alterations (11,118). However, our findings propose that the core symptoms might stem from microglia overactivation, a phenomenon potentially triggered not only by dopamine deficit but also by other mechanisms. This insight sheds light on the intricate etiology of ADHD, aligning with existing evidence indicating its association with inflammatory conditions such as allergic or atopic diseases (119,120). Furthermore, it underscores the observation that a subset of patients fails to respond to current methylphenidate or atomoxetine treatments.

Hence, our study offers a plausible rationale for the diverse presentations of the disorder, whether rooted in genetic dopamine dysfunction or environmental inflammatory factors. Consequently, effective patient stratification becomes imperative for tailoring appropriate treatments and minimizing adverse effects.

## Acknowledgements

We thank University Jaume I Central Services; *Servicios de Experimentación Animal (SEA)* for animal care and Confocal microscopy services. Illustrations were created with *BioRender* software.

## Disclosures

### Author Contributions

Conceptualization, AMSP, MMB, SSS. Software, MMB, SSS; validation, MMB, SSS, AMSP; formal analysis, MMB, AMSP; investigation, MMB, AMSP; resources, MMB, AMSP; data curation, MMB, SSS, AMSP; writing—original draft preparation, MMB, AMSP; writing—review and editing, AMSP, SSS, MMB, NK, ML, MRL; visualization, AMSP; supervision, AMSP; project administration, AMSP; funding acquisition, AMSP, NK, ML, SSS. All authors have read and agreed to the published version of the manuscript.

### Institutional Review Board Statement

The study was approved by the Ethics Committee of the University Jaume I. Protocol code 2020/VSC/PEA/0099); the date of approval 5/June/2020.

### Data Availability Statement

Data available upon request.

---

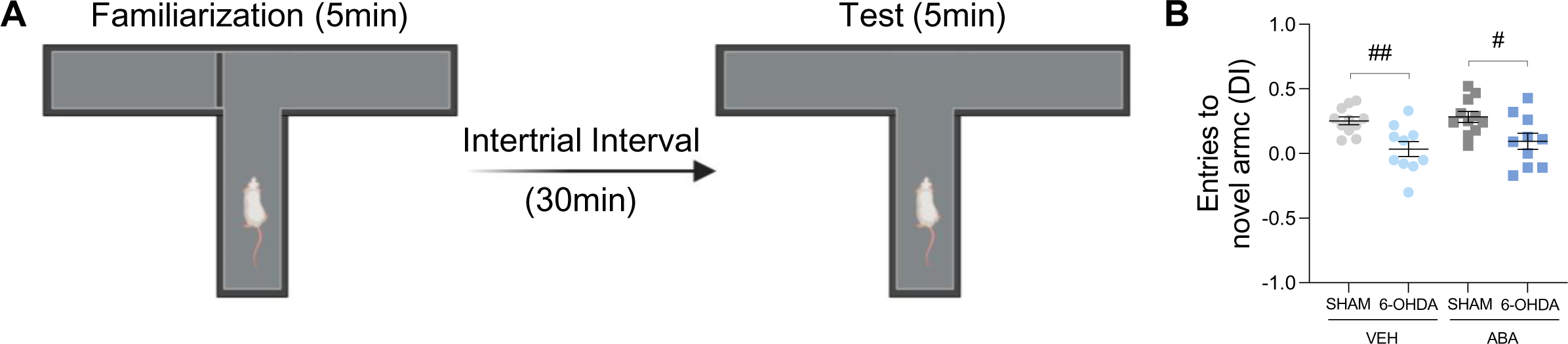

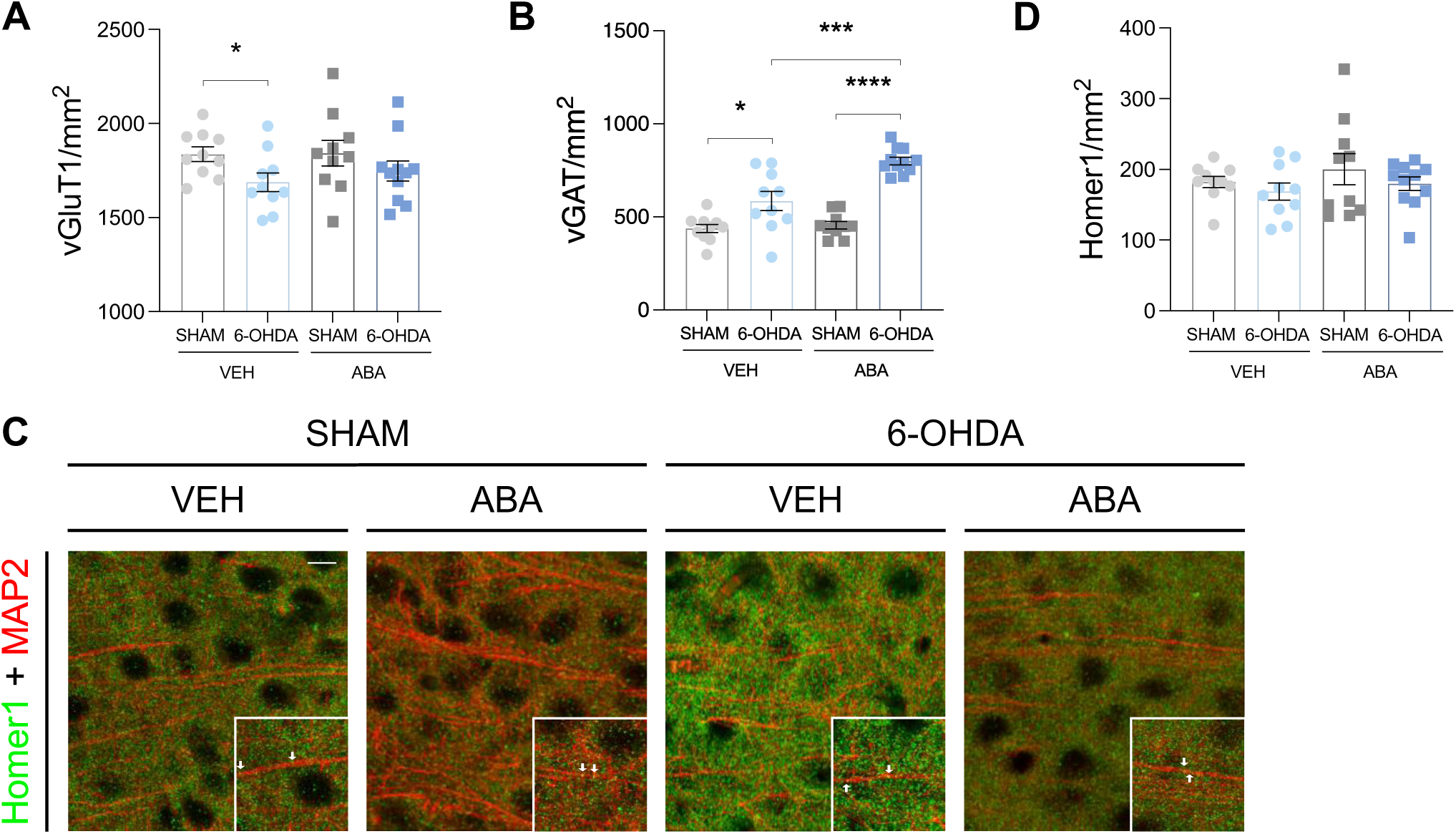

**Supplementary data. Tables 1-4.**

**Means ± SEM** comparing lesion effect, and treatment effect (Unpaired student t-test).

**Table 1.** Effect of neonatal 6-OHDA lesion observed at P21.

| Behavior | SHAM | 6-OHDA | P value<br>(Unpaired Student t-test) |
| --- | --- | --- | --- |
| Locomotor activity. Distance (cm) | 4095 ± 124 | 5730 ± 305.4 | p < 0.0001 **** (t = 4.30, df = 49) |
| Locomotor activity. Velocity (cm/sec) | 6.80 ± 0.20 | 9.70 ± 0.47 | p < 0.0001 **** (t = 4.94 df = 49) |
| Impulsivity. Number of entries (%) | 39.22 ± 2.33 | 39.84 ± 3.18 | p = 0.8755 (t = 0.16 df = 41) |
| Impulsivity. Time (%) | 18.13 ± 1.72 | 34.10 ± 5.20 | p = 0.0050 ** (t = 2.97 df = 41) |
| Anxiety. Number of entries center | 15.45 ± 1.43 | 26.75 ± 2.53 | p < 0.0001 **** (t = 5.225 df = 40) |
| Anxiety. Time in center (sec) | 9.19 ± 1.04 | 14.13 ± 1.41 | p = 0.0072 ** (t = 2.84, df = 39) |
| Anxiety. Latency to first crossing (sec) | 48.25 ± 7.22 | 37.65 ± 6.60 | p = 0.286 (t = 1.08, df = 39) |
| Mechanical Stimuli. Filament (grams) | 1.69 ± 0.10 | 0.47 ± 0.03 | p < 0.0001 **** (t = 11.67 df = 19) |
| Social Interaction. Time (%) | 52.14 ± 2.635 | 65.23 ± 3.98 | p = 0.0080 ** (t = 2.79 df = 40) |
| NOR (d-index) | 0.025 ± 0.05 | -0.15 ± 0.05 | p = 0.0192 * (t = 2.42 df = 48) |

**Table 2.**
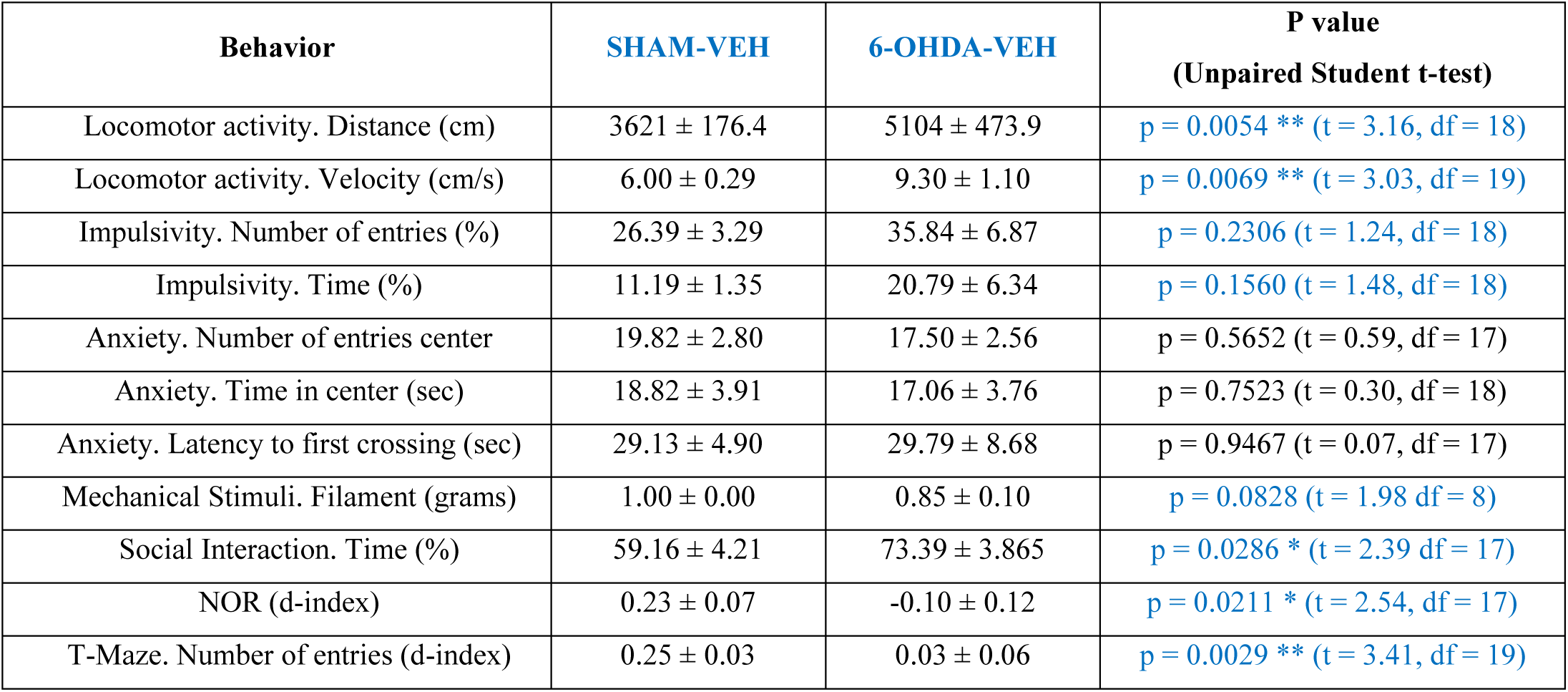
Effect of neonatal 6-OHDA lesion observed at P90. Untreated.

**Table 3.**
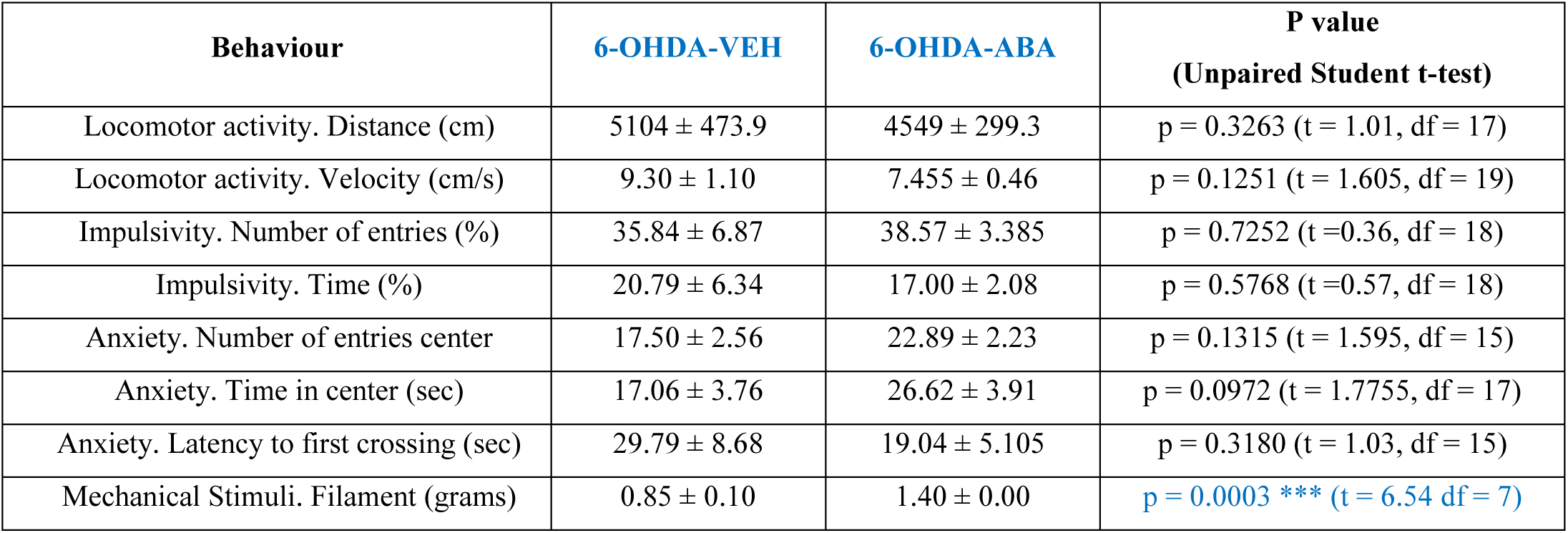
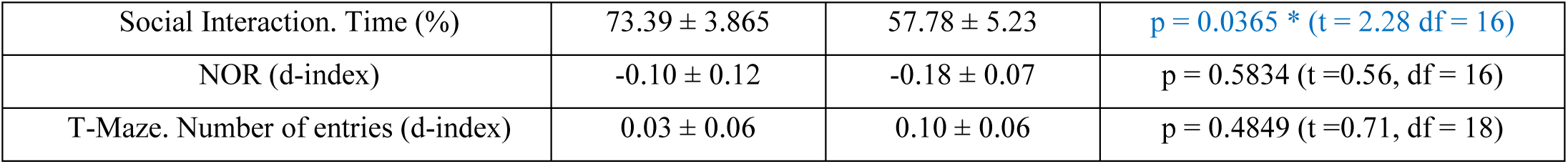
Effect of ABA treatment in neonatal 6-OHDA lesion at P90.

| Behaviour | 6-OHDA-VEH | 6-OHDA-ABA | P value<br>(Unpaired Student t-test) |
| --- | --- | --- | --- |
| Locomotor activity. Distance (cm) | 5104 ± 473.9 | 4549 ± 299.3 | p = 0.3263 (t = 1.01, df = 17) |
| Locomotor activity. Velocity (cm/s) | 9.30 ± 1.10 | 7.455 ± 0.46 | p = 0.1251 (t = 1.605, df = 19) |
| Impulsivity. Number of entries (%) | 35.84 ± 6.87 | 38.57 ± 3.385 | p = 0.7252 (t = 0.36, df = 18) |
| Impulsivity. Time (%) | 20.79 ± 6.34 | 17.00 ± 2.08 | p = 0.5768 (t = 0.57, df = 18) |
| Anxiety. Number of entries center | 17.50 ± 2.56 | 22.89 ± 2.23 | p = 0.1315 (t = 1.595, df = 15) |
| Anxiety. Time in center (sec) | 17.06 ± 3.76 | 26.62 ± 3.91 | p = 0.0972 (t = 1.7755, df = 17) |
| Anxiety. Latency to first crossing (sec) | 29.79 ± 8.68 | 19.04 ± 5.105 | p = 0.3180 (t = 1.03, df = 15) |
| Mechanical Stimuli. Filament (grams) | 0.85 ± 0.10 | 1.40 ± 0.00 | p = 0.0003 *** (t = 6.54 df = 7) |
| Social Interaction. Time (%) | 73.39 ± 3.865 | 57.78 ± 5.23 | p = 0.0365 * (t = 2.28 df = 16) |
| NOR (d-index) | -0.10 ± 0.12 | -0.18 ± 0.07 | p = 0.5834 (t = 0.56, df = 16) |
| T-Maze. Number of entries (d-index) | 0.03 ± 0.06 | 0.10 ± 0.06 | p = 0.4849 (t = 0.71, df = 18) |

**Table 4.**
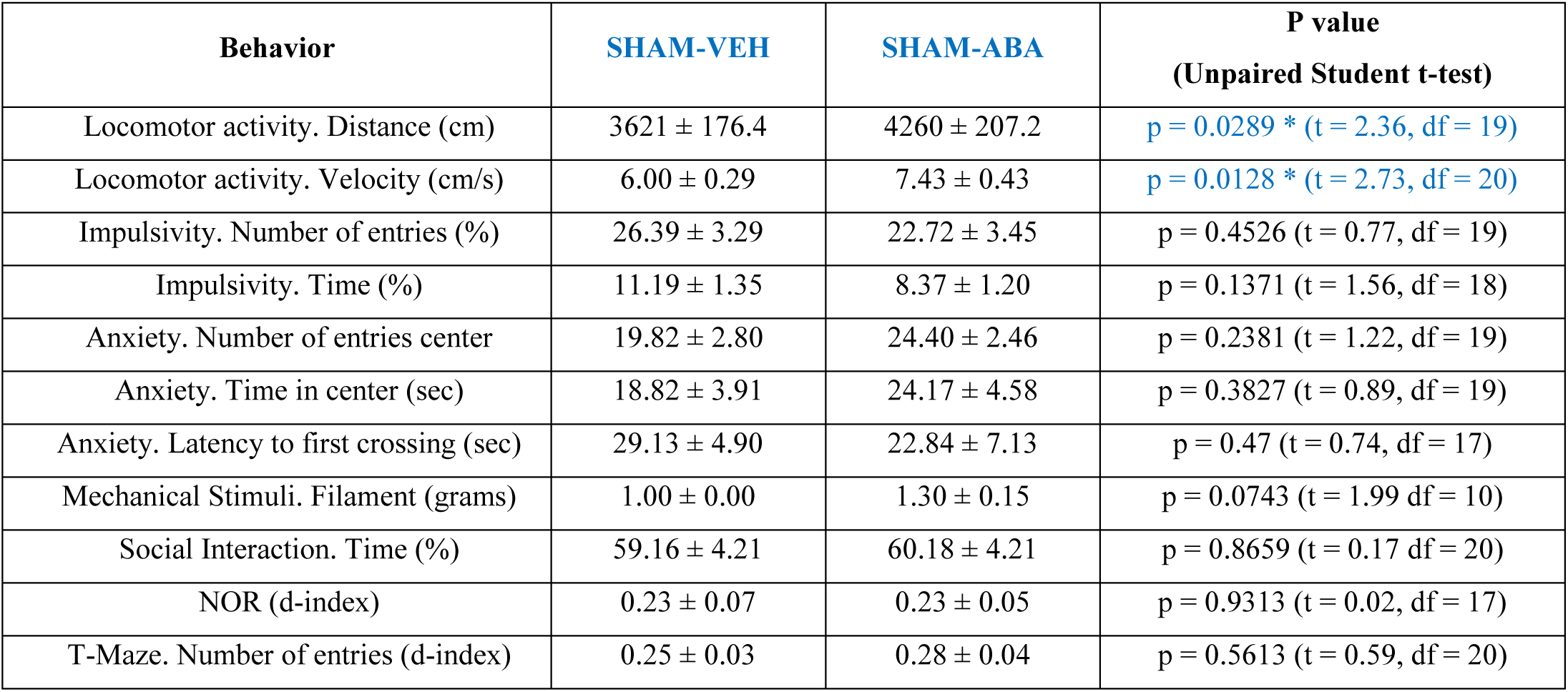
Effect of ABA treatment in Sham mice at P90.

